# The role of muscle stem cells and fibro-adipogenic progenitors in female pelvic floor muscle regeneration following birth injury

**DOI:** 10.1101/2021.07.30.454534

**Authors:** Francesca Boscolo Sesillo, Varsha Rajesh, Michelle Wong, Pamela Duran, Brittni Baynes, Louise C. Laurent, Karen L. Christman, Alessandra Sacco, Marianna Alperin

**Author notes:** Corresponding author: Marianna Alperin, MD, MS, University of California, San Diego, 9500 Gilman Drive, La Jolla, CA 92093-0863. These authors equally contributed to this work (Francesca Boscolo Sesillo, Varsha Rajesh). **Author contributions:** Conceptualization, F.B.S, L.C.L., and M.A.; Methodology, F.B.S., and M.A.; Formal Analysis, F.B.S., and M.A.; Investigation, F.B.S., V.R., M.W., P.D., B.B.; Resources, K.L.C., M.A.; Writing – Original Draft, F.B.S., and V.R.; Writing – Review & Editing, F.B.S., P.D., L.C.L., K.L.C., A.S., M.A., Visualization, F.B.S.; Supervision, F.B.S., and M.A.; Project Administration, F.B.S.; Funding Acquisition, M.A.

## Abstract

Pelvic floor muscle (PFM) injury during childbirth is a key risk factor for subsequent pelvic floor disorders that affect millions of women worldwide. Muscle stem cells (MuSCs) play a central role in the regeneration of injured skeletal muscles, where they activate, proliferate, and differentiate to assure myogenesis needed for muscle recovery. For robust regenerative function, MuSCs require the support of fibro-adipogenic progenitors (FAPs) and immune cells. To elucidate the role of MuSCs, FAPs, and immune infiltrate in female PFM regeneration, we used radiation to perturb the system and followed PFM recovery in a simulated birth injury (SBI) rat model. Non-irradiated and irradiated rats were euthanized at 3,7, 10, and 28 days after SBI; PFMs were harvested and prepared for immunohistochemistry. Cross sectional area (CSA) of all PFM myofibers 28 days after injury in irradiated animals was significantly lower relative to non-irradiated injured controls, indicating impairment of PFM recovery. Following SBI in non-irradiated animals, the number of MuSCs and FAPs expanded significantly at 7 and 3 days after injury, respectively; this expansion did not occur in irradiated animals at the same time points. CSA of embryonic myosin heavy chain (eMyHC, marker of newly regenerated myofibers) positive fibers was also significantly smaller following SBI in irradiated muscles compared to PFMs from non-irradiated injured controls at 7 days. Our results demonstrate that loss of function and decreased expansion of MuSCs and FAPs associated with irradiation results in impaired PFM recovery, signifying essential roles for MuSCs and FAPs in the regenerative process of female PFMs after birth injury. These findings can inform the identification of novel preventative and therapeutic targets and the development of new treatments for PFM dysfunction and associated pelvic floor disorders.

## Introduction

Skeletal muscle regeneration consists of a tightly orchestrated series of events that involve multiple cell types. Muscle stem cells (MuSCs) are muscle resident cells, which play a major role in the regenerative process (Lepper et al., 2011; Relaix and Zammit, 2012). During muscle regeneration, MuSCs, which are quiescent in homeostatic conditions, become activated, proliferate, and eventually differentiate and fuse with existing myofibers or form new ones (Dumont et al., 2015). The contributions of fibro-adipogenic progenitors (FAPs) and immune cells are essential to achieve efficient regeneration (Arnold et al., 2007; Joe et al., 2010). FAPs are muscle resident mesenchymal cells located in the interstitial space between myofibers (Joe *et al*., 2010; Uezumi et al., 2010). Similar to MuSCs, they start proliferating after muscle injury, supporting MuSCs activation and expansion through cytokine secretion (Biferali et al., 2019; Joe *et al*., 2010). Immune cells are recruited to the site of injury as early as 1 hour after muscle damage occurs (Chazaud et al., 2009; Tidball, 2005). Neutrophils that are the “first responders” secrete pro-inflammatory cytokines, inducing the recruitment of monocytes and macrophages, to the site of injury (Tidball, 2005). Macrophages first differentiate into pro-inflammatory (M1) and subsequently anti-inflammatory (M2) phenotypes in response to changes in the environmental cues and cytokine milieu (Chazaud *et al*., 2009). Interactions among these different cell types are essential for proper muscle regeneration (Biferali *et al*., 2019; Dort et al., 2019).

Female pelvic floor muscles (PFMs) span the pelvic outlet to support pelvic and abdominal viscera, aid in urinary and fecal continence, and enable sexual function. In humans, PFMs include the levator ani complex (the medial (puborectalis) and lateral (pubococcygeus) portions of pubovisceralis, and iliococcygeus) and the posterior coccygeus muscle (Gowda and Bordoni, 2020). PFM dysfunction is one of the leading contributors to the development of pelvic floor disorders (PFDs) that include pelvic organ prolapse, and urinary and fecal incontinence (Hallock and Handa, 2016). These morbid and costly conditions negatively impact quality of life of close to a quarter of the U.S. female population (Hallock and Handa, 2016). Maternal trauma to the PFMs during vaginal delivery due to eccentric contractions (natural childbirth) or passive stretch (childbirth under regional anesthesia) is the leading risk factor for PFM dysfunction; however, not all women develop this condition consequent to childbirth (Hallock and Handa, 2016). One of the contributing factors associated with the development of PFDs is increased maternal age (Hallock and Handa, 2016; Memon and Handa, 2013; Urbankova et al., 2019). Aging has been also associated with loss of MuSC and FAP function, leading to the impairment in muscle regeneration (Hwang and Brack, 2018; Lukjanenko et al., 2019; Moratal et al., 2019). Thus, it is possible that the disparity in PFM dysfunction consequent to parturition is, at least in part, due to the differential endogenous potential for PFM regeneration. Despite a well-established correlation between PFM birth injury and maternal age at the time of delivery with subsequent development of PFDs (DeLancey et al., 2003; DeLancey et al., 2007; Memon and Handa, 2013), little research has been done to investigate PFM regeneration following vaginal childbirth. Thus, the above served as the main impetus for the current study.

The majority of existing studies focused on cellular changes during muscle regeneration are performed in murine hind limb muscles subjected to either myotoxic (cardiotoxin, notexin), chemical (barium chloride), acute denervation, or cryo-injury (Gayraud-Morel et al., 2009; Hardy et al., 2016; Musarò, 2014). Moreover, these studies are conducted primarily in male animal models, leaving the female muscle regenerative biology highly understudied. Significant sexual dimorphisms exist in skeletal muscles. At the transcriptional level, numerous genes involved in regulation of muscle mass are differentially expressed in male vs. female muscles, resulting in abundant differences, including fiber phenotype and contractility (Haizlip et al., 2015; Welle et al., 2008). Thus, studies focused on female muscle regeneration in general, and pelvic skeletal muscles specifically, are urgently needed to bridge the existing scientific disparity. Furthermore, despite dramatic prevalence of PFDs that disproportionately affect parous women, to our knowledge, there are no published studies evaluating the role of MuSCs, FAPs, or immune infiltrate in postpartum PFM regeneration.

Given the ethical and technical limitations associated with the use of human subjects for the studies of PFMs, we utilized the rat model, previously validated for the investigations of human PFMs and the impact of birth injury on pelvic soft tissues (Alperin et al., 2014; Catanzarite et al., 2018; Hoyte et al., 2008; Lien et al., 2004). The rat PFM complex is analogous to humans, consisting of the pubocaudalis (PCa) and iliocaudalis (ICa) muscles that together comprise the rat levator ani, and the coccygeus (C) muscle (Alperin *et al*., 2014). Employing the simulated birth injury (SBI) rat model, we aimed to assess the contribution of MuSCs and FAPs during PFM regeneration following birth injury. We hypothesized that, similarly to other skeletal muscles, MuSCs and FAPs play major roles in female PFM constructive remodeling after birth injury. To explicitly test our hypothesis, we relied on irradiation, which has been shown to induce breaks in the DNA structure analogous to the phenotype observed in cellular senescence and aging (Borrego-Soto et al., 2015; Marcoux et al., 2013; Ness et al., 2015), to perturb MuSC and FAP function.

## Results

### Pelvic floor muscle regeneration following simulated birth injury is impaired after whole muscle irradiation

To assess the regenerative potential of PFMs and determine cellular events necessary for muscle recovery following SBI, we employed irradiation with a single 20Gy dose (Figure 1A), which is in the range shown to comprise MuSC function in limb muscles. Quiescent and low proliferating cells are less affected than highly proliferating cells, thus, to perturb these cells functionality, a higher irradiation dose is required (Borrego-Soto *et al*., 2015). Indeed, the MuSC function has been shown to be compromised when 18-25 Gy are applied to the muscle (Vahidi Ferdousi et al., 2014). SBI was performed via vaginal balloon distention that replicates circumferential and downward strains imposed on PFMs during parturition (Alperin et al., 2010). First, we sought to determine the impact of stem and progenitor cellular impairment on muscle regeneration by comparing the histomorphology of PCa 3, 7, and 28 days post injury in irradiated and non-irradiated animals (Figure 1B–C). PCa was selected as a starting point because it sustains the largest strain during delivery compared to the other component of the rat levator ani complex, analogous to the pubovisceralis portion of human levator ani (Catanzarite *et al*., 2018; Hoyte *et al*., 2008; Lien *et al*., 2004; Sindhwani et al., 2017). Our findings are consistent with the published data, with all PCa muscles examined demonstrating large injured areas, whereas only two-thirds of the ICa samples exhibited signs of muscle damage. Approximately 20% of the C muscle was spared from the impact of SBI. The extent of injury was also evaluated indirectly, through the assessment of the embryonic myosin heavy chain (eMyHC) expression in the non-irradiated muscles 7 days post SBI, and was found to be the highest in PCa, followed by C, and then ICa, consistent with the histomorphological muscle appearance (Figure S1A-B).

**Figure 1.**
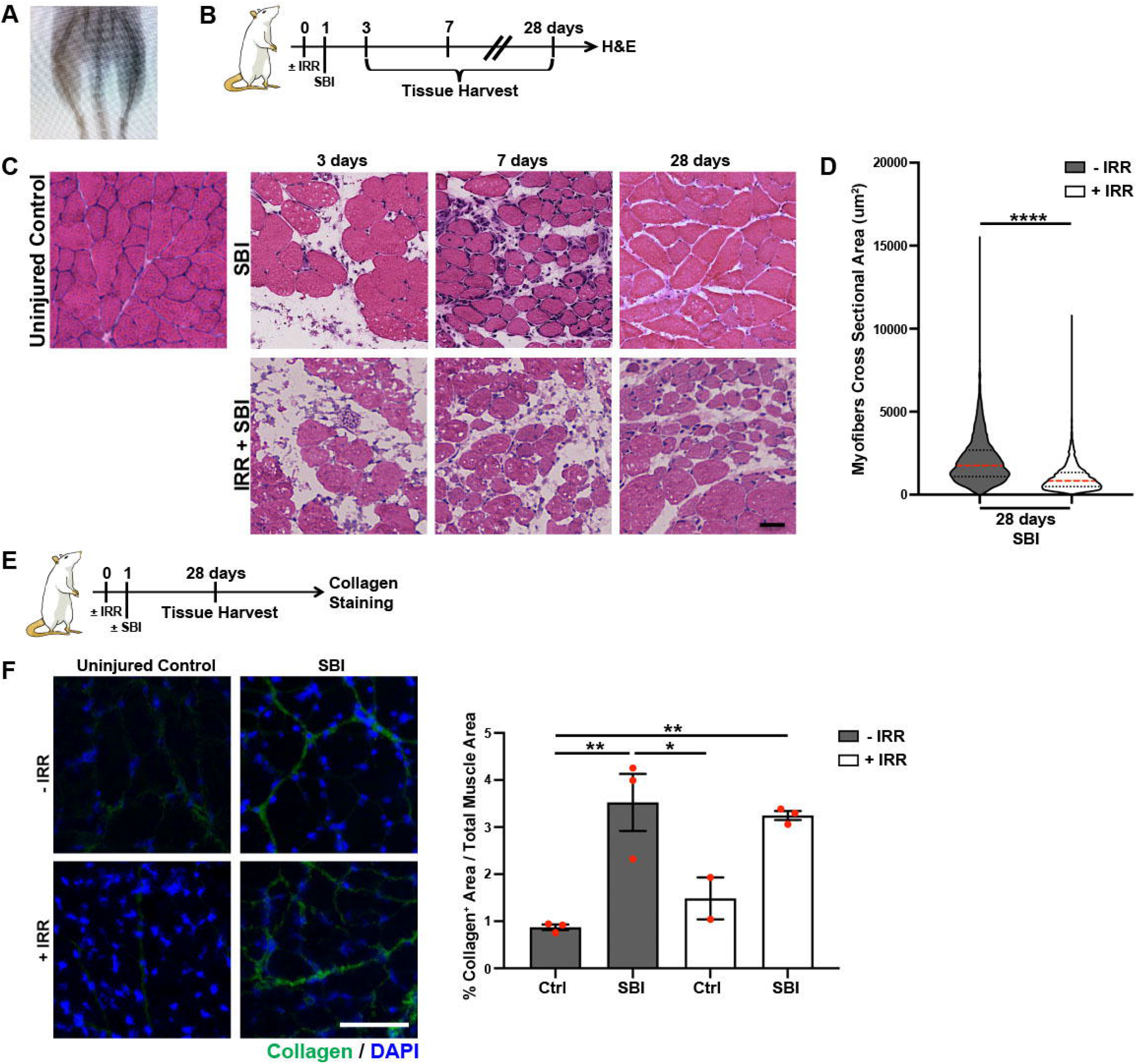
The impact of simulated birth injury on the morphometric properties of pubocaudalis (PCa) in non-irradiated and irradiated rat model. (**A**) X-ray image of the irradiated area. (**B**) Schematic representation of the experimental design for C. (**C**) H&E staining of PCa at 3, 7, and 10 days after injury (scale bar: 50μm). (**D**) Violin plots representing distribution of the myofiber size at 28-day time point. ****: p-value< 0.0001; Mann-Whitney test. n=3 animals. (**E**) Schematic representation of the experimental design for F. (**F**) On the left, representative immunofluorescent images of collagen staining (scale bar: 100μ m). On the right, quantification of the collagen area fraction. Red dots represent individual measurements; error bar represent SEM. *: p-value < 0.05; **: p-value < 0.01. One-way ANOVA with Tukey’s post-hoc. n=3 animals.

The extent of injury in PCa from the irradiated and non-irradiated groups was comparable, as evident from the large interstitial spaces 3 days post SBI (Figure 1C). As expected, regenerating myofibers, identified by centralized nuclei, were present in non-irradiated animals 7 days post injury. On the other hand, animals irradiated before injury did not show signs of muscle regeneration at this time point (Figure 1C). One month after injury, the differences between the two groups were even more pronounced. While PCa morphology in non-irradiated injured animals did not fully return to the unperturbed state, PCa in irradiated injured animals exhibited significantly smaller myofibers and persistent large interstitial spaces despite a 4-week recovery period, demonstrating substantial impairment in regeneration (Figure 1C–D). Taken together, the above indicates that irradiation effectively impairs PFMs regenerative ability and thus can be efficiently employed as a tool to identify the major players in PFMs’ recovery following birth injury. Interestingly, the myofiber size of ICa and C did not differ between non-irradiated and irradiated animals, suggesting that complete regeneration might have occurred in these muscles or that the initial amount of injury in these muscles was different than PCa (Figure S1C-D).

With respect to extracellular matrix (ECM), we found that the total content of the PCa collagen, a major constituent of the intramuscular ECM identified via immunostaining (Figure 1E), was not altered by irradiation in uninjured muscles when compared to non-irradiated controls (Figure 1F). Consistent with the previous studies conducted in our lab (Duran et al., 2021), we observed significant increase in the intramuscular collagen content in non-irradiated injured PFMs compared to uninjured controls (Figure 1F). Irradiation prior to SBI did not further increase fibrotic degeneration of injured PCa relative to non-irradiated counterparts (Figure 1F).

### Irradiation impairs pelvic floor muscle stem cell proliferative capacity and fibro-adipogenic progenitor expansion in response to simulated birth injury

Irradiation is known to negatively impact the function of dividing cells and thus, we hypothesized that it affects MuSC and FAP functionality following birth injury, leading to the long-term phenotype described above. To explicitly test this hypothesis, we first compared the response of PFM stem cells to birth injury in non-irradiated and animals irradiated 1 day before SBI (Figure 2A, Figure S2A). As an initial step, we confirmed that the MuSC reservoir is not altered by irradiation in the otherwise unperturbed PFMs by employing Pax7 to identify and quantify MuSCs *in situ*. Pax7^+^ cell number did not differ one day after irradiation in any of the 3 individual PFMs (PCa, ICa, and C) compared to non-irradiated uninjured animals (Figure 2B, Figure S2B-2C, Table S1). The importance of the above is that at the time of SBI, the starting MuSC reservoir was not different between non-irradiated and irradiated animals.

**Figure 2.**
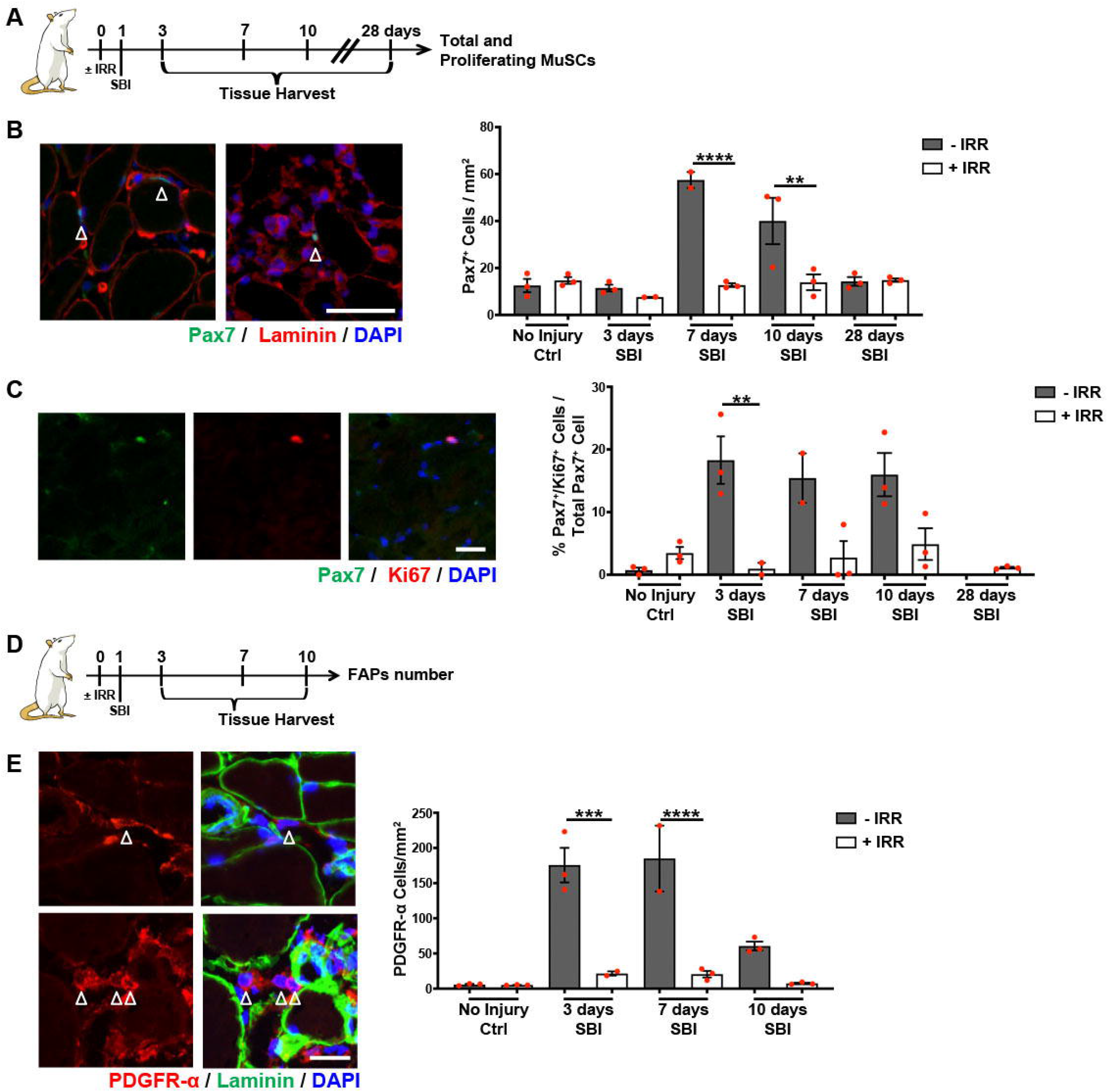
Muscle stem cells (MuSCs) and fibro-adipogenic progenitors (FAPs) behavior in regenerating pubocaudalis (PCa) muscle in non-irradiated and irradiated animals. (**A**) Schematic representation of the experimental design for B and C. (**B, C,** and **D**) Red dots represent single measurements; error bar represent SEM. (**B**) On the left, representative immunofluorescent images of Pax7/laminin staining of non-irradiated injured (left) and irradiated injured (right) PFMs 7 days following simulated birth injury. White arrows indicate MuSCs (scale bar: 50μm). On the right, quantification of MuSC number per mm^2^. **: p-value < 0.01; ****: p-value< 0.0001; One-way ANOVA with Tukey’s post-hoc. Complete Statistics in Table S1. n=3 animals. (**C**) On the left, representative immunofluorescent images of Pax7/Ki67 staining (scale bar: 25μm). On the right, quantification of Pax7/Ki67 double positive cells. *: p-value < 0.05; **: p-value < 0.01; One-way ANOVA with Tukey’s post-hoc. n=3 (**D**) Schematic representation of the experimental design for E. (**E**) On the left, representative immunofluorescent images of PDGFR-α/laminin staining. Top two images are of uninjured non-irradiated pubocaudalis; bottom two images are of non-irradiated pubocaudalis 3 days after simulated birth injury. White arrows indicate FAPs (scale bar: 25 μ m). On the right, quantification of FAP number per mm^2^. Red dots represent single measurements; error bar represent SEM. **: p-value < 0.01; ***: p-value < 0.001; ****: p-value< 0.0001; One-way ANOVA with Tukey’s post-hoc. n=3 animals.

While the number of PCa MuSCs was similar in irradiated and non-irradiated animals 3 days after SBI (Figure 2B), significant increase in the number of MuSCs was observed at 7 and 10 days following birth injury in non-irradiated animals, consistent with a previous study performed by our group (Duran *et al*., 2021). In contrast, this upsurge was not recapitulated in the irradiated group (Figure 2B, Table S1). The number of MuSCs returned to basal levels 28 days after injury in the non-irradiated group (Figure 2B, Table S1). Similar results were observed for C, while MuSC number of ICa, which experiences lower strains during SBI (Catanzarite *et al*., 2018), did not differ significantly between non-irradiated and irradiated animals at any time point during the recovery period (Figure S2B-2C). These findings suggest that irradiation perturbs MuSC activation and/or proliferation and that this effect is modulated by the magnitude of strain and related muscle injury.

To determine potential cause of the quantitative differences in MuSCs following SBI between irradiated and non-irradiated groups, we went on to assess MuSC proliferation. Employing Ki67 as a marker of cell proliferation, we identified and quantified proliferating MuSCs in the regenerating PFMs. The number of proliferating MuSCs did not differ between irradiated and non-irradiated uninjured animals in any of the 3 individual PFMs (Figure 2C, Figure S2D-2E, Table S2). However, consistent with the overall quantitative differences post-SBI, the proportion of proliferating MuSCs in PCa and C amplified only in the non-irradiated group, with no increase in MuSC proliferation observed in the irradiated animals. In contrast, no significant difference between groups was observed in ICa. PFM stem cell proliferation returned to baseline at 28 days after SBI (Figure 2C, Figure S2D-2E, Table S2). These results demonstrate, for the first time, that irradiation of PFMs negatively impacts MuSC proliferative ability during regeneration following birth injury, similarly to what has been observed in hind limb muscles (Vahidi Ferdousi *et al*., 2014).

In addition to MuSCs, FAPs that aid in MuSC activation and proliferation are another important player in proper skeletal muscle regeneration. To our knowledge, direct proof of the negative effect of irradiation on FAP activation does not exist in the currently published literature. Thus, we first asked whether the function of these cells is affected by irradiation. Subsequently, we went on to determine the role of FAPs in PFM recovery following birth injury. We performed immunohistochemistry using anti-PDGFR-α antibody to identify and quantify FAPs in uninjured and injured (3, 7, and 10 days post SBI) PFMs of irradiated and non-irradiated animals (Figure 2D). We did not observe any differences in the PCa FAP number in non-injured animals with or without irradiation (Figure 2E, Table S3). In response to injury, FAP number significantly increased in PCa of non-irradiated animals 3 and 7 days post SBI, whereas this increase was not observed in irradiated injured animals at either time point (Figure 2E, Table S3). The FAP number was similar to the baseline level 10 days after injury (Figure 2E, Table S3). Findings in C were consistent with those in PCa; however, similar to the response of ICa MuSCs, the FAP number in this component of the PFM complex did not differ significantly between irradiated and non-irradiated injured animals (Figure S2F-2H). Importantly, these results suggest that the ability of PFM FAPs to expand after SBI is impaired by irradiation, likely contributing to the inadequate PFM regeneration in response to birth injury.

### Blood vessels and immune response of the pelvic floor muscles after birth injury are not affected by irradiation

Proper muscle regeneration also requires a temporally regulated immune response, where circulating leukocytes are recruited from the vasculature to the site of injury to clear the debris and aid in activation and differentiation of MuSCs and FAPs (Dort *et al*., 2019). To determine whether irradiation affects intramuscular blood vessels and, therefore, potentially impairs the access of immune cells to PFMs, we examined arterioles at 8 and 28 days after irradiation using an antibody against α-smooth muscle actin (α-SMA), a smooth muscle cell marker (Figure 3A, Figure S3A). Irradiation alone did not affect arteriolar size or density at either time point (Figure 3B, Figure S3B-3C). In addition, we did not observe any difference in the number of infiltrating cells in irradiated vs non-irradiated muscles acutely after SBI (Figure S3D).

**Figure 3.**
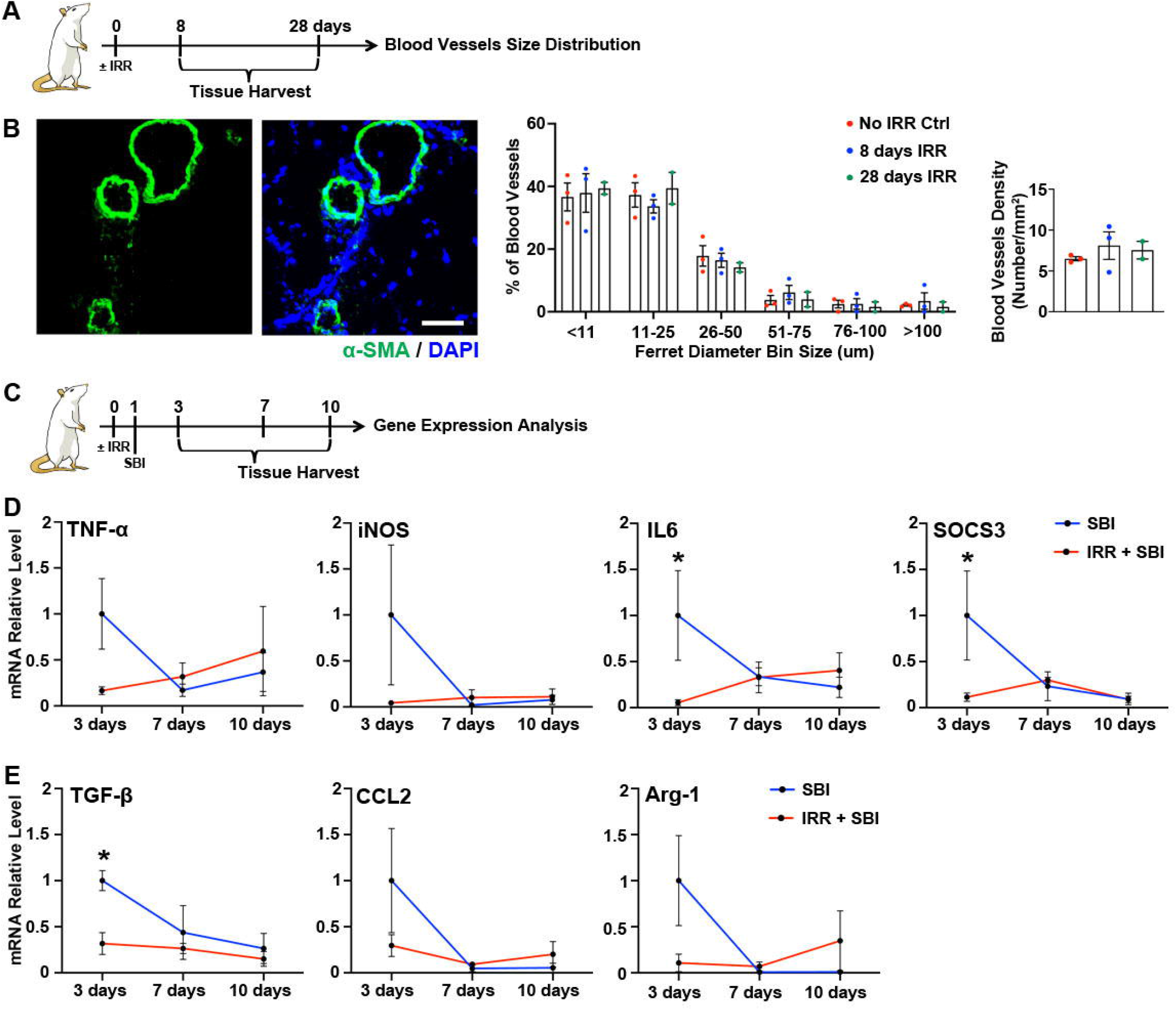
Blood vessels and immune infiltrate in regenerating pubocaudalis (PCa) muscle in non-irradiated and irradiated animals. (**A**) Schematic representation of the experimental design for B. (**B**) On the left, representative immunofluorescent images of α-SMA staining (scale bar: 50μm). In the center, bar graph representing the distribution of blood vessel ferret diameters. On the right, bar graph representing blood vessel density. Colored dots represent single measurements; error bar represent SEM. One-way ANOVA with Tukey’s post-hoc. n=3 animals. (**C**) Schematic representation of the experimental design for D and E. (**D**) qPCR analysis of pro-inflammatory genes. *: p-value < 0.05; Two-way ANOVA with Dunnett’s multiple comparison test. n=3 animals. (**E**) qPCR analysis of anti-inflammatory genes. *: p-value < 0.05; Two-way ANOVA with Dunnett’s multiple comparison test. n=3 animals.

Given that irradiation did not impact infiltrating cells at the site of muscle injury-regeneration quantitatively, we then asked whether the immune response mounted by irradiated animals after birth injury was comparable to that of non-irradiated group. To address this question, we designed a panel of genes associated with pro- and anti-inflammatory responses of injured skeletal muscles and compared their expression between irradiated and non-irradiated animals in the whole muscle at 3, 7, and 10 days post SBI (Figure 3C, Figure S3E). Out of all genes examined (Table S4), IL6 and SOCS3 (pro-inflammatory markers) and TGFB1 (anti-inflammatory marker) were upregulated in PCa 3 days after injury in non-irradiated compared to irradiated animals (Figure 3D–3E). None of the genes showed a significant increase in expression at any time point in irradiated injured animals (Figure 3D–3E). When the same genes were analyzed in C and ICa, we observed a different pattern: SOCS3 was upregulated in both muscles in irradiated samples 7 days after injury (Figure S3F-3G). In C, we also observed upregulation of iNOS (pro-inflammatory marker) at 3 days and IL6 at 7 days after SBI in irradiated animals (Figure 3G). To determine whether the differential expression observed in the individual components of the PFM complex is due to the variable effect of radiation versus disparate muscle response to SBI, we assessed the expression of the same genes in the tibialis anterior (TA), the hind limb muscle that was subjected to irradiation but not to birth injury. We did not observe any difference in the gene expression of the TA muscle when irradiated samples were compared to non-irradiated ones (Figure S3H). Taken together, these results indicate that irradiation does not affect the ability of PFMs to mount an immune response, with observed differences attributable to the known variable extent of injury in the individual components of the PFM complex (Catanzarite *et al*., 2018).

### Early muscle regeneration is impaired in irradiated regenerating PFM

Given the reduction in MuSC proliferative ability, the lack of FAPs expansion, and the clear phenotypic differences observed in irradiated versus non-irradiated PFMs 4 weeks after birth injury, we went on to further explore the events during acute and subacute recovery period. Previously, we have demonstrated the presence of embryonic myosin heavy chain (eMyHC) positive fibers in PCa as early as 3 days post SBI in non-irradiated animals REF HERE. Using the same anti-eMyHC antibody, we assessed myofiber regeneration at 3, 7, and 10 days post injury in the irradiated group (Figure 4A, Figure S4A). In contrast to the observations in non-irradiated animals, eMyHC^+^ fibers were absent in irradiated injured animals at 3 days post SBI (Figure 4B, left panel). At 7 days, the size of eMyHC^+^ fibers was significantly smaller in irradiated compared to non-irradiated rats (Figure 4B, right panel). The difference in eMyHC^+^ fiber size between irradiated and non-irradiated animals was no longer observed at 10 days post injury (Figure 4B right panel). The above findings indicate that irradiated animals have a delayed start in the regenerative process, whereas non-irradiated animals with intact MuSC and FAP function start the regenerative process as early as 3 days after injury. Similar trends were observed in the ICa and C muscles (Figure S4B-4C). Taken together, these results indicate that deficient MuSC proliferation and lack of FAP expansion impair female PFMs regeneration following birth injury.

**Figure 4.**
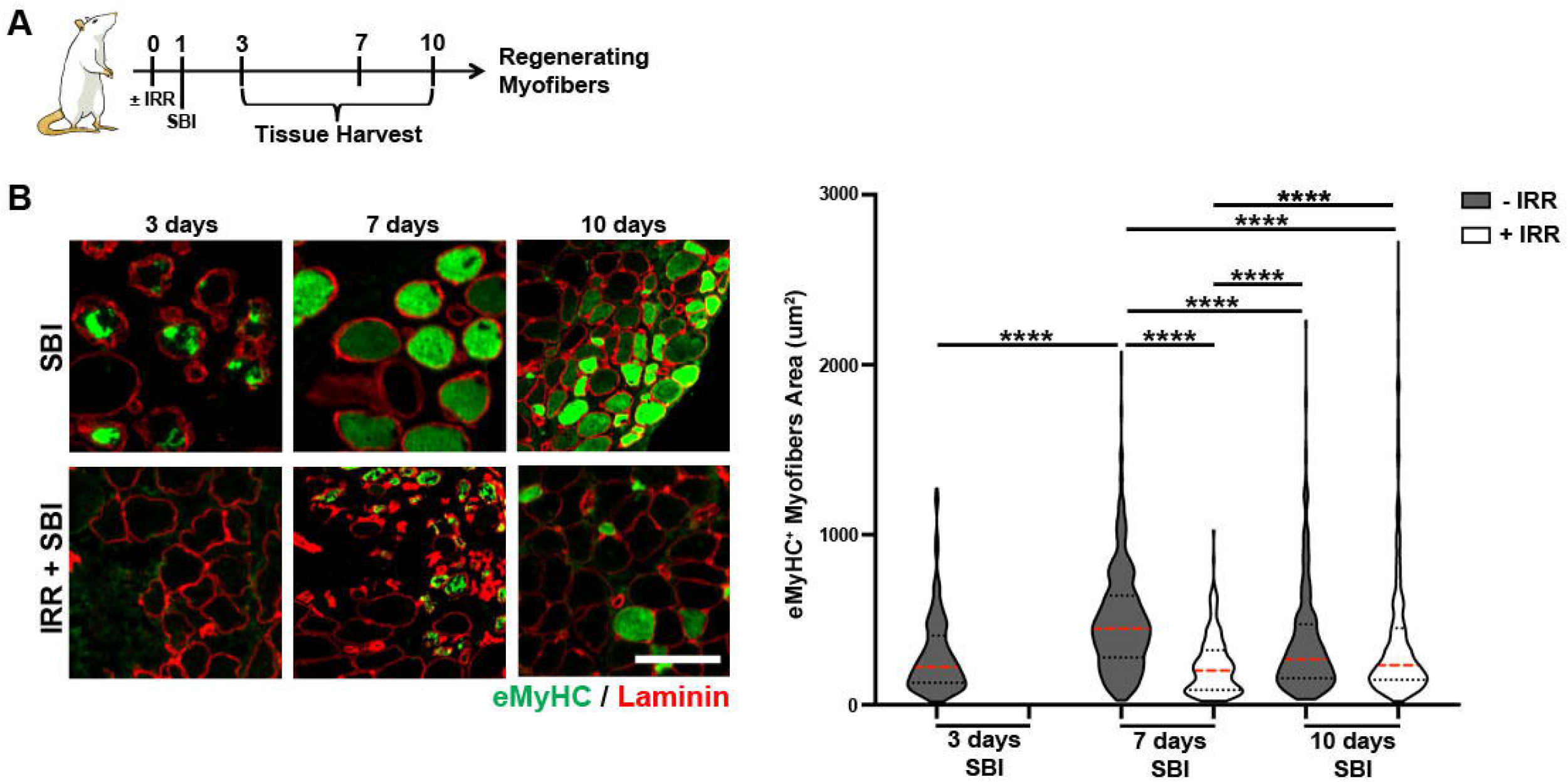
Early pubocaudalis (PCa) muscle regeneration in non-irradiated and irradiated animals. (**A**) Schematic representation of the experimental design for B. (**B**) On the left, representative immunofluorescent images of eMyHC/laminin staining (scale bar: 100μm). On the right, violin plots representation of eMyHC^+^ fiber area. ****: p-value< 0.0001; Kruskal-Wallist with Dunn’s multiple comparison test. n=3 animals.

## Discussion

Our novel study examines the functional role of the major players (MuSCs and FAPs) in female PFMs regeneration after birth injury. Employing a validated simulated birth injury rat model and irradiation as a tool to perturb the main contributors to skeletal muscle regeneration, we show that PFM cross sectional area is substantially reduced in irradiated compared to non-irradiated animals 4 weeks after birth injury. Accumulation of collagen content was observed in injured compared to uninjured animals, as previously described (Duran *et al*., 2021), but was not exacerbated further by irradiation in the injured animals. We demonstrate that in injured non-irradiated animals FAPs and MuSCs increase in number at 3 and 7 days following SBI, leading to the activation of the regenerative process. In contrast, MuSCs and FAPs in irradiated PFMs do not expand during the first stages of muscle regeneration following birth injury. Moreover, the regenerating muscle fibers do not appear in irradiated PFMs until day 7 post SBI. Immune infiltrate does not appear to be impacted by irradiation. Taken together, these results show that the coordinated activity of MuSCs and FAPs is necessary to ensure proper PFMs recovery after injury and that delay in the regenerative process leads to a substantial reduction in myofibers cross sectional area 4 weeks after birth injury.

In the current study, we observed a variable impact of birth injury on MuSCs and FAPs in the individual components of the rat PFM complex. Specifically, in response to SBI, MuSC number and proliferation increased in PCa and C, but not in ICa. Likewise, FAP number increased acutely after injury only in the PCa and C muscles. Notably, we observed large injury-regeneration areas in PCa and C, assessed by eMyHC expression, whereas the extent of injury was smaller and more variable in ICa. These observations are consistent with our previous findings of the differential PFMs’ response to mechanical birth injury, owing to the variable strain magnitude imposed on the individual components of the PFM complex during SBI (Catanzarite *et al*., 2018). Specifically, Catanzarite *et al*., showed that PCa and C were significantly more affected by SBI, displaying dramatic myofiber stretch, and sarcomere hyperelongation and misalignment, while ICa was less affected (Catanzarite *et al*., 2018). Our discoveries in the rat PFMs are consistent with the findings of computer modeling of human parturition and clinical radiological studies, which show that the iliococcygeus (analogous to ICa) portion of the levator ani muscle experiences less strain than pubococcygeus (analogous to PCa) during fetal delivery (Hoyte *et al*., 2008; Lien *et al*., 2004; Sindhwani *et al*., 2017). Unfortunately, the coccygeus has been omitted from the majority of the human studies this far. Here, we show for the first time that parturition-related strains of variable magnitudes imposed on each PFM during SBI correspond to disparate cellular responses to birth injury. Indeed, ICa, which experiences the smallest parturition-associated strains, demonstrates substantially reduced cellular responses to SBI compared to PCa and C. Our findings are also consistent with the investigations conducted in hind limb muscles, where MuSC and FAP increase in number after injury, while cells in uninjured muscles do not change quantitatively over time (Joe *et al*., 2010; Lemos et al., 2015; Murphy et al., 2011; Wang et al., 2014b).

The existence of multiple transgenic mouse models makes the study of the relative contribution to regeneration of MuSCs, FAPs, and immune cells relatively easy in the murine model (Brigitte et al., 2010; Lepper *et al*., 2011; von Maltzahn et al., 2013; Wang et al., 2014a; Wosczyna et al., 2019). Given the absence of rat genetic models, we employed irradiation as a tool to determine the role of various cellular populations in female PFM regeneration. It has been previously shown that the MuSC compartment in hind limb muscles is affected by irradiation. Indeed, radiation induces accumulation of DNA damage in MuSCs that, in turn, inhibits cellular capacity to proliferate and differentiate into mature myofibers after injury (Boldrin et al., 2012; Gulati, 1987; Vahidi Ferdousi *et al*., 2014). Moreover, MuSC self-renewal ability is also reduced by irradiation (Vahidi Ferdousi *et al*., 2014). Consistent with previous observations in appendicular muscles, we demonstrate that the impairment of PFM stem cells’ ability to enter the cell cycle and proliferate results in a delay in the formation of regenerating myofibers and a reduction in their cross-sectional area during early regeneration.

The effect of irradiation on FAPs has not been previously described (D’Souza et al., 2019; Doreste et al., 2020). Nevertheless, it has been hypothesized that because FAPs become activated in response to muscle injury, their function could be compromised by irradiation (Doreste *et al*., 2020; Joe *et al*., 2010). Here, we provide the first evidence that the function of FAPs is directly affected by irradiation. Given the major role that FAPs play in regulating activation and proliferation of MuSCs, their loss of function in our model system may contribute to the observed defect in tissue repair. FAPs are also a major source of intramuscular collagen, thus, their impaired ability to activate in response to injury might partially account for the absence of a higher fibrotic reaction in irradiated relative to non-irradiated injured PFMs 4 weeks after birth injury (Uezumi et al., 2011). However, we cannot exclude the possibility that pathological deposition of collagen, originating from other sources, might accumulate in irradiated PFMs at a later time point leading to worsening fibrosis, as it has been previously shown in both human and animal studies (Hsu et al., 1998; Zhou et al., 2018).

We further show that irradiation does not affect either the size or density of arterioles up to one month after irradiation. Similarly to our finding, a single high dose irradiation of dystrophic animals has been shown not to affect endothelial cells up to 28 days after treatment (Boldrin *et al*., 2012). In addition, another study shows that immediately after a single dose irradiation, blood flow in skeletal muscles is increased, and this increase persists up to 4 week after irradiation, suggesting that blood vessel function is not altered negatively after irradiation (Song et al., 1983).

Studies aiming to determine the effect of irradiation on inflammatory response in regenerating muscles are also limited (Doreste *et al*., 2020; Robertson et al., 1992). In a chronic inflammatory muscle environment, such as muscular dystrophy, irradiation has been shown to increase the innate immune response, inducing a favorable environment for the engraftment of donor derived MuSCs (Doreste *et al*., 2020). Instead, when a healthy muscle was injured 24 hours after local irradiation, no difference in the amount of infiltrating cells was observed during the early regenerative process (Robertson *et al*., 1992). Consistent with this study that better reflects our experimental conditions, we did not observe significant overall differences in either pro- or anti-inflammatory PFMs’ response to birth injury in irradiated vs non-irradiated animals.

Interestingly, despite employing a high-dose pelvic gamma radiation that negatively affected proliferation of pelvic muscles’ MuSCs and FAPs, we still observed a certain level of muscle regeneration. Previous studies have demonstrated that a subpopulation of MuSCs is capable of forming fully differentiated myofibers upon transplantation despite 18 Gy irradiation (Boldrin *et al*., 2012). It is also known that this MuSC subpopulation is activated in response to injury (Heslop et al., 2000). Moreover, it has been recently reported that radiation-resistant MuSCs have enhanced Pax3 expression and can expand *in vivo* after injury despite irradiation, contributing to muscle regeneration (Scaramozza et al., 2019). Pax3 expression is necessary for protection of cells from external stresses, preventing premature cellular activation and cell death through activation of G-alert state, characterized by accelerated cell cycle entry (Der Vartanian et al., 2019). Exploring whether incomplete loss of the PFMs’ regenerative capacity in the irradiated injured group is due to the presence of radiation-resistant MuSCs is a fruitful avenue for the future studies.

Irradiation is a well-known cause of cellular senescence, a cell state defined by irreversible cell cycle arrest, apoptotic resistance, and altered gene expression (Campisi and d’Adda di Fagagna, 2007; Chen et al., 2019). Published studies demonstrate that cancer survivors rapidly accumulate senescent cells and prematurely develop aging traits after undergoing radiation therapy (Marcoux *et al*., 2013; Ness *et al*., 2015). MuSCs in aged or diseased conditions have also been shown to become senescent and negatively affect skeletal muscle homeostasis and regeneration (Childs et al., 2015; García-Prat et al., 2016; He and Sharpless, 2017; Sacco et al., 2010; Sousa-Victor et al., 2014). To date, almost nothing is known about the effects of cellular senescence and aging on the female PFMs (Rieger et al., 2021). In the future, the irradiation model used in this study could be used as a tool to induce cellular senescence and premature PFM aging to further the knowledge in the field of female pelvic medicine.

In conclusion, we demonstrate that PFM constructive regeneration following birth injury is dependent on MuSC and FAP ability to become activated and proliferate. Therefore, these cell populations could be used as actionable therapeutic targets to enhance endogenous PFM repair response in women at high risk of maternal pelvic floor injury (operative vaginal delivery, macrosomia, prolonged second stage of labor) or with potentially diminished ability for adequate PFM recovery following birth injury (older maternal age, extensive muscle injury, previous substantial muscle injuries, immune compromise).

## Methods

### Animals

Female 3-month old Sprague-Dawley rats (Envigo, Indianapolis, IN) were randomly divided in 4 study groups: (1) unperturbed controls; (2) animals subjected to simulated birth injury (SBI) only; (3) animals undergoing irradiation only; and (4) animals subjected to irradiation and SBI. The rats were allowed to recover for either 3, 7, 10, or 28 days post simulated birth injury (SBI), at which point they were euthanized by CO_2_ inhalation followed by bilateral thoracotomy (N=3/group/time point). Bilateral coccygeus (C), iliocaudalis (ICa), pubocaudalis (PCa), and non-pelvic control muscles - tibialis anterior (TA) were harvested immediately following euthanasia. All procedures were approved by the University of California San Diego Institutional Animal Care and Use Committee.

### Simulated birth injury (SBI)

SBI was performed using an established vaginal distention protocol, as described in Alperin et. al 2010 (Alperin *et al*., 2010). Briefly, the rats were anesthetized with 2.5% isoflurane with oxygen for the duration of the procedure. A 12-French transurethral balloon catheter (Bard Medical, Covington, GA) with the tip cut off was inserted into the vagina with 130 grams weight attached to the end of the catheter. The balloon was inflated to 5 ml and left in place for 2 hours, after which it was pulled through the introitus to simulate the circumferential and downward distention associated with fetal crowning and parturition.

### Irradiation

Animals were irradiated at UCSD animal facility with the Small Animal IGRT Platform (SmART+) from Precision X-Ray Irradiation using SmART Advanced Treatment Planning software. Pelvic floor and hind limb muscles were irradiated with a single dose of 20 Gy (225 kV, 20mA with a treatment filter).

### RNA Isolation and Quantitative PCR

Harvested PCa, C, ICa, and TA were homogenized in liquid nitrogen for RNA isolation. RNA was isolated using the RNeasy Mini Kit (QIAGEN, 74106) following the manufacturer’s protocol, and quantified using QIAxpert (QIAGEN, 9002340). cDNA was synthesized using the SuperScript™ IV First-Strand Synthesis System (ThermoFisher Scientific, 18091050), and RT-qPCR was performed using Power SYBR™ Green PCR Master Mix (ThermoFisher Scientific, 4367659), on CFX96 Touch™ Real-Time PCR Detection System (Bio-Rad, 1855195) with 1 ng cDNA and primers listed in Table S4. The control gene used was 60S acidic ribosomal protein P0 (Rplp0), and Ct values for all genes were divided by the Rplp0 Ct value to obtain relative gene expression.

Out of all the genes in Table S4, IL-4 and IL-10 were excluded from the analysis since their expression in the majority of the samples was above 35 Cts.

### Immunostaining

Snap frozen muscles were sectioned into 10 μm thick slices and fixed for immunostaining in either acetone (Fisher Scientific, A16P-4), 4% PFA (Fisher Scientific, 04042) or 2% PFA, depending on the primary antibody used. eMyHC and α-SMA slides where incubated in sequence with PBS (phosphate buffered saline) and blocking buffer (20% normal goat serum (Gemini Bio-products, 100-109) + 0.3% Triton X-100 (Sigma-Aldrich, X100-500ML) in PBS) before overnight incubation with primary antibody. For Pax7 and Ki67 staining, slides were washed in PBS and then placed in antigen unmasking solution (Vector Laboratories, H-3301-250) for antigen retrieval. Sections were washed with PBS and incubated in blocking buffer before adding the primary antibody. Slides stained with PDGFR-α were first washed with PBS, then incubated with a 0.5% Triton X-100 solution and finally with casein blocking buffer (0.25% casein (Sigma, C3400-500G) in PBS) before overnight incubation with primary antibody. Slides stained for collagen I were first washed and then incubated in in blocking buffer (10% goat serum, 0.3% Triton x-100, 1% BSA (Gemini Bio-Products, 700-100P) in PBS) before overnight incubation with primary antibody. Slides were washed in PBS and then incubated with secondary antibody in blocking buffer for 1 hour. To identify cell nuclei, slides were incubated in DAPI (ThermoFisher Scientific, 62248, 1:1000) in PBS for 10 min.

Primary antibodies used were: rabbit anti-laminin (Sigma, L9393, 1:200), mouse anti-embryonic myosin heavy chain (Developmental Studies Hybridoma Bank (DSHB), F1.652, 1:200), mouse anti-Pax7 (Developmental Studies Hybridoma Bank (DSHB), Pax7-c, DSHB, 1:100), and rabbit anti-Ki67 (Abcam, ab15580, 1:100), goat anti-PDGFR-α (R&D Systems, AF1062, 1:100), α-SMA (Cell Signaling, 19245S, 1:100), collagen I (Invitrogen, PA5-95137, 1:200). Secondary antibodies used were: Alexa Fluor 488 goat anti-mouse IgG (Invitrogen, A21121, 1:200 for eMyHC and 1:250 for Pax7), and Alexa Fluor 546 goat anti-rabbit IgG (Invitrogen, A11035, 1:500 for laminin and 1:250 for Ki67), Alexa Fluor 546 donkey anti-goat IgG (Invitrogen, A-11056, 1:250), Alexa Fluor 488 donkey anti-rabbit (Invitorgen, A-21206, 1:250).

### Hematoxylin and Eosin staining

Snap frozen tissue sections were first rinsed in water, then immerse in Harris’s Hematoxylin for 3 minutes, followed by water rinses and 30 dips in lithium carbonate solution. Slides where then dip in 1% eosin solution in acetic acid, and de-hydrated in ethanol. Slides where finally dipped in Citri-Soly until they became clear and finally protected with a coverslip.

### Imaging

Immunofluorescence imaging was carried out using the LEICA fluorescent microscope (LEICA AF6000 Modular System). Slides stained for fibrosis were imaged with a Leica Aperio ScanScope® CS2.

### Quantification

#### Embryonic myosin heavy chain

For each muscle (PCa, ICa, or C) taken from each animal, 1 tissue section containing the largest observable area of injury from either or both the left or right side of the muscle were chosen for analysis. Full images of each section were captured at 10X, and the area of each eMyHC^+^ muscle fiber was measured. Myofibers cross sectional area was assessed using ImageJ 1.51s and ImageJ64. Images were not modified before quantification.

#### Pax7/ki67

For each rat muscle section, 3 images were taken at 20X of each tissue section, from both the right and left sides. The number of Pax7^+^ cells, and Pax7^+^/ki67^+^ cells were counted in each image. The number of Pax7^+^ cells was calculated per unit area, while Pax7^+^/ki67^+^ double positive cells were expresses as percentage. Unmodified images were analyzed using ImageJ 1.51s and Adobe Photoshop CS4.

#### Pdgfr-α

For each uninjured muscle section, 3 images were taken at 20X of each tissue section, from both the right and left sides. The number of PDGFR- α^+^ cells, and cells were counted in each image. For injured muscle, only the sections containing an injury were imaged at 20X and used for quantification. The number of PDGFR-α^+^ cells was calculated per unit area. Unmodified images were analyzed using ImageJ 1.51s and Adobe Photoshop CS4.

#### Collagen

One slide per muscle sample, containing 10-14 tissue sections, was analyzed employing ImageJ 1.51s. Specifically, all sections were outlined with the freehand selections tool (edges of the muscle section, nerves, and large blood vessels were excluded from the analysis), converted into binary images, and quantified as percentage area occupied by collagen staining.

### Statistical Analysis

Data were analyzed using GraphPad Prism v8.0, San Diego, CA. Distribution was assessed using Kolmogorov-Smirnov test for normality. Data that followed a parametric distribution, such as collagen content, MuSC number and percentage of proliferating cells, FAP number, and blood vessels size and density were compared using one-analysis of variance (ANOVA) followed by Tukey’s post hoc pairwise comparisons, when indicated. Non-parametrically distributed variables, such as fiber area, were analyzed by Kruskal-Wallis test followed by Dunn’s pairwise comparisons (for eMyHC^+^ fibers) or with Mann-Whitney test (for 28 days after injury time point). Gene expression data were analyzed using a two-way ANOVA with Dunnett’s post-hoc test with a significance set to p<0.05.

## Supplementary Figure Legends

***Figure S1 – Iliocaudalis (ICa) and coccygeus (C) response to injury in non-irradiated and irradiated animals*** (**A**) Representative immunofluorescent images of eMyHC/laminin staining in PCa, ICa, and C 7 days after injury. (B) Quantification of percentage of eMyHC^+^ areas over the total muscle section area. Colored dots represent single measurements; error bar represent SEM. One-way ANOVA with Tukey’s post-hoc. n=2-3

***Figure S2 – Muscle stem cells (MuSCs) and fibro-adipogenic progenitors (FAPs) behavior in regenerating Iliocaudalis (ICa) and coccygeus (C) muscles in non-irradiated and irradiated animals*** (**A**) Schematic representation of the experimental design for B to E. (**B-E** and **G-H**) Red dots represent single measurements; error bar represent SEM. (**B**) Bar graph representing the quantification of MuSCs number per mm^2^ in ICa muscle. One-way ANOVA with Tukey’s post-hoc. n=3 (**C**) Bar graph representing the quantification of MuSCs number per mm^2^ in C muscle. *: p-value < 0.05; **: p-value < 0.01; ***: p-value < 0.001; One-way ANOVA with Tukey’s post-hoc. n=3 (**D**) Bar graph representing the quantification of Pax7/Ki67 double positive cells in ICa muscle. One-way ANOVA with Tukey’s post-hoc. n=3 (**E**) Bar graph representing the quantification of Pax7/Ki67 double positive cells in C muscle. *: p-value < 0.05; **: p-value < 0.01; ***: p-value < 0.001; ****: p-value< 0.0001; One-way ANOVA with Tukey’s post-hoc. n=3. If the number of red dots within one sample is <3, it means that one of the analyzed samples did not show injured area and thus could not be quantified. (**D**) Schematic representation of the experimental design for G and H. (**G**) Bar graph representing the quantification of FAPs number per mm^2^ in ICa muscle. (**H**) Bar graph representing the quantification of FAPs number per mm^2^ in C muscle. *: p-value < 0.05; **: p-value < 0.01; One-way ANOVA with Tukey’s post-hoc. n=3

***Figure S3 – Blood vessels and immune infiltrate in regenerating Iliocaudalis (ICa) and coccygeus (C) muscle in non-irradiated and irradiated animals*** (**A**) Schematic representation of the experimental design for B and C. (**B-C**) Colored dots represent single measurements; error bar represent SEM. (**B**) On the left, bar graph representing the distribution of blood vessel ferret diameters in ICa muscle. On the right, bar graph representing blood vessel density. One-way ANOVA with Tukey’s post-hoc. (**C**) On the left, bar graph representing the distribution of blood vessel ferret diameters in C muscle. On the right, bar graph representing blood vessel density. One-way ANOVA with Tukey’s post-hoc. (**D**) Representative immunofluorescent images of laminin and DAPI 3 days after injury (scale bar: 50 μm). (**E**) Schematic representation of the experimental design for panels F to H (**F**) On the top, qPCR analysis of pro-inflammatory genes for ICa muscle. On the bottom qPCR analysis of anti-inflammatory genes for ICa muscle. Two-way ANOVA with Dunnett’s multiple comparison test. (**G**) On the top, qPCR analysis of pro-inflammatory genes for C muscle. On the bottom qPCR analysis of anti-inflammatory genes for C muscle. *: p-value < 0.05; **: p-value < 0.01; Two-way ANOVA with Dunnett’s multiple comparison test. (**H**) On the top, qPCR analysis of pro-inflammatory genes for TA muscle. On the bottom qPCR analysis of anti-inflammatory genes for TA muscle. Two-way ANOVA with Dunnett’s multiple comparison test.

**Figure S4 – *Early Iliocaudalis (ICa) and coccygeus (C) muscles regeneration in non-irradiated and irradiated animals*** (**A**) Schematic representation of the experimental design for B and C. (**B**) Violin plots representation of eMyHC^+^ fiber area for ICa muscle. ***: p-value < 0.001; ****: p-value< 0.0001; Kruskal-Wallist with Dunn’s multiple comparison test. (**C**) Violin plots representation of eMyHC^+^ fiber area for C muscle. **: p-value < 0.01; ****: p-value< 0.0001; Kruskal-Wallist with Dunn’s multiple comparison test.

**Table S1.**

| **Tukey's multiple comparisons test** | **Significance** |
| --- | --- |
| No Injury Ctrl -IRR vs 7 days SBI -IRR | **** |
| No Injury Ctrl -IRR vs 10 days SBI -IRR | ** |
| No Injury Ctrl +IRR vs 7 days SBI -IRR | **** |
| No Injury Ctrl +IRR vs 10 days SBI -IRR | ** |
| 3 days SBI -IRR vs 7 days SBI | **** |
| 3 days SBI -IRR vs 10 days SBI | ** |
| 3 days SBI +IRR vs 7 days SBI | **** |
| 3 days SBI +IRR vs 10 days SBI | ** |
| 7 days SBI -IRR vs. 7 days SBI + IRR | **** |
| 7 days SBI -IRR vs 10 days SBI + IRR | **** |
| 7 days SBI -IRR vs 28 days SBI | **** |
| 7 days SBI -IRR vs 28 days SBI +IRR | **** |
| 7 days SBI +IRR vs 10 days SBI | ** |
| 10 days SBI -IRR vs 10 days SBI + IRR | ** |
| 10 days SBI -IRR vs 28 days SBI | ** |
| 10 days SBI -IRR vs 28 days SBI +IRR | ** |

**Table S2.**

| **Tukey's multiple comparisons test** | **Significance** |
| --- | --- |
| No Injury Ctrl -IRR vs 3 days SBI -IRR | ** |
| No Injury Ctrl -IRR vs 7 days SBI -IRR | * |
| No Injury Ctrl -IRR vs 10 days SBI -IRR | ** |
| No Injury Ctrl +IRR vs 3 days SBI -IRR | * |
| No Injury Ctrl +IRR vs 10 days SBI -IRR | * |
| 3 days SBI -IRR vs 7 days SBI +IRR | ** |
| 3 days SBI -IRR vs 10 days SBI +IRR | * |
| 3 days SBI -IRR vs 28 days SBI +IRR | ** |
| 3 days SBI +IRR vs 10 days SBI -IRR | * |
| 7 days SBI -IRR vs 28 days SBI +IRR | * |
| 7 days SBI +IRR vs 10 days SBI -IRR | * |
| 10 days SBI -IRR vs 28 days SBI +IRR | * |

**Table S3.**
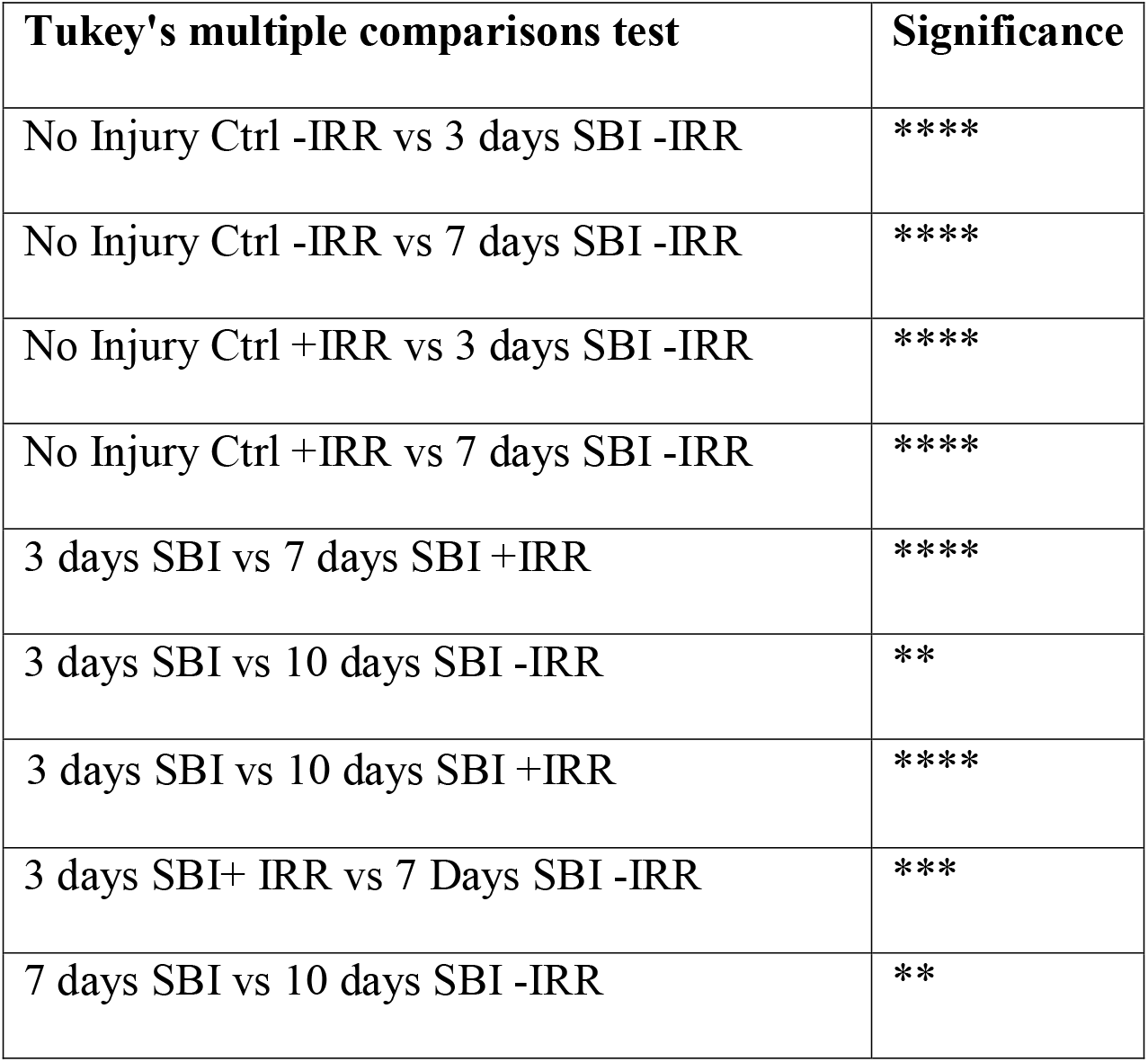
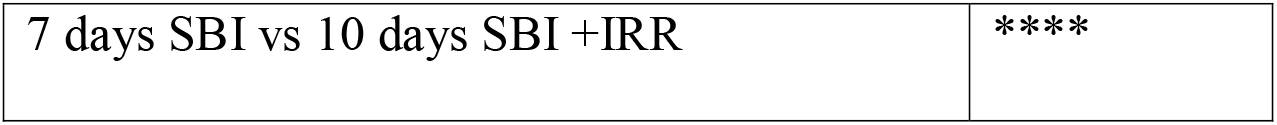

**Table S4.**
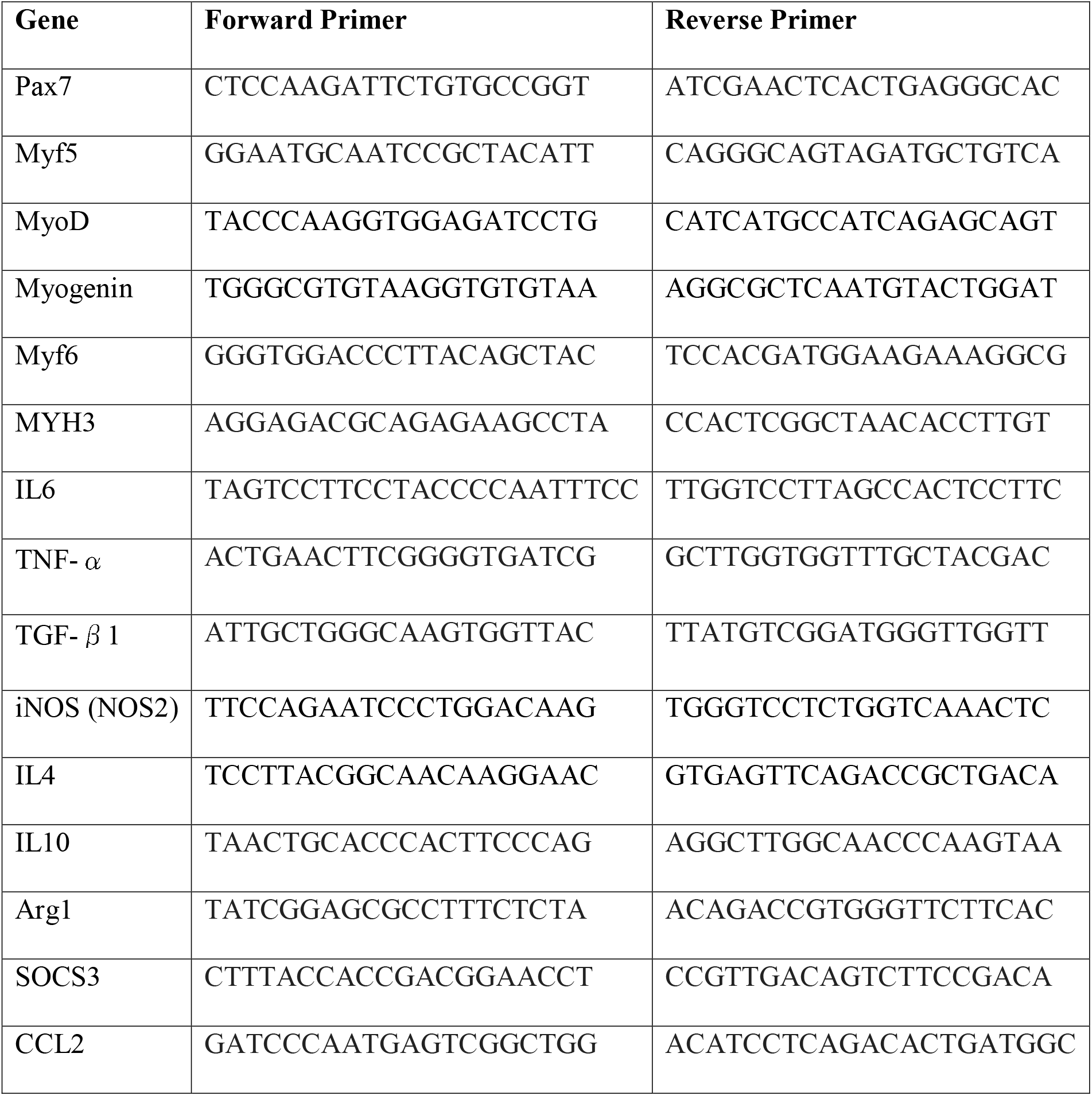

## Acknowledgement

UCSD Animal Care Program Technical Services

