## Supplementary figures and images for "The role of muscle stem cells and fibro-adipogenic progenitors in female pelvic floor muscle regeneration following birth injury"

### Supplemental Figure 1

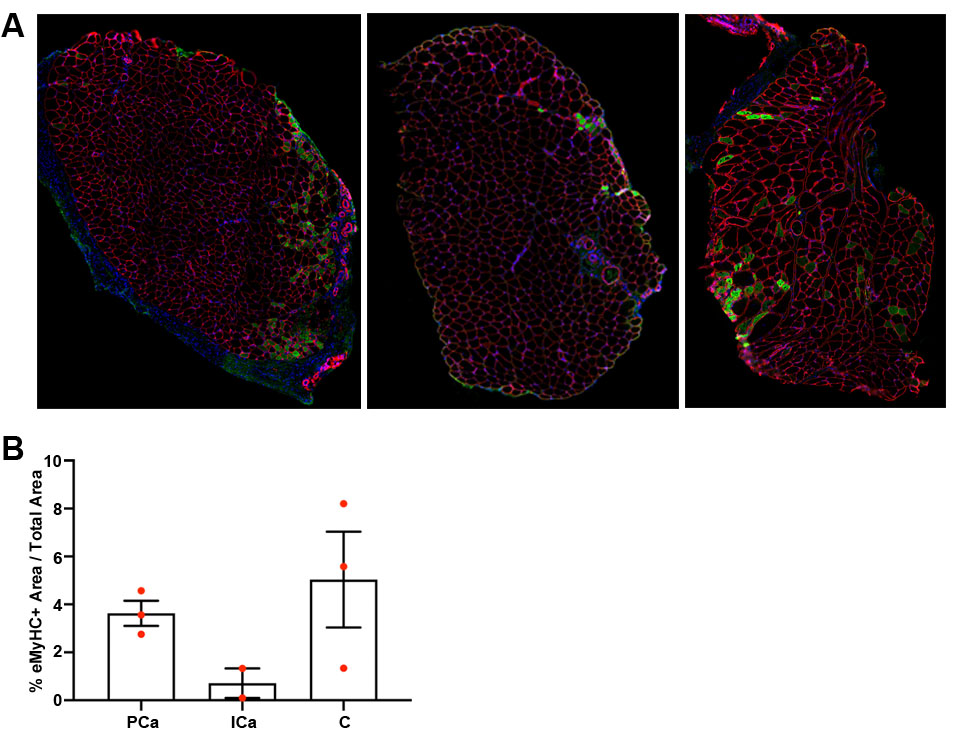

### Supplemental Figure 2

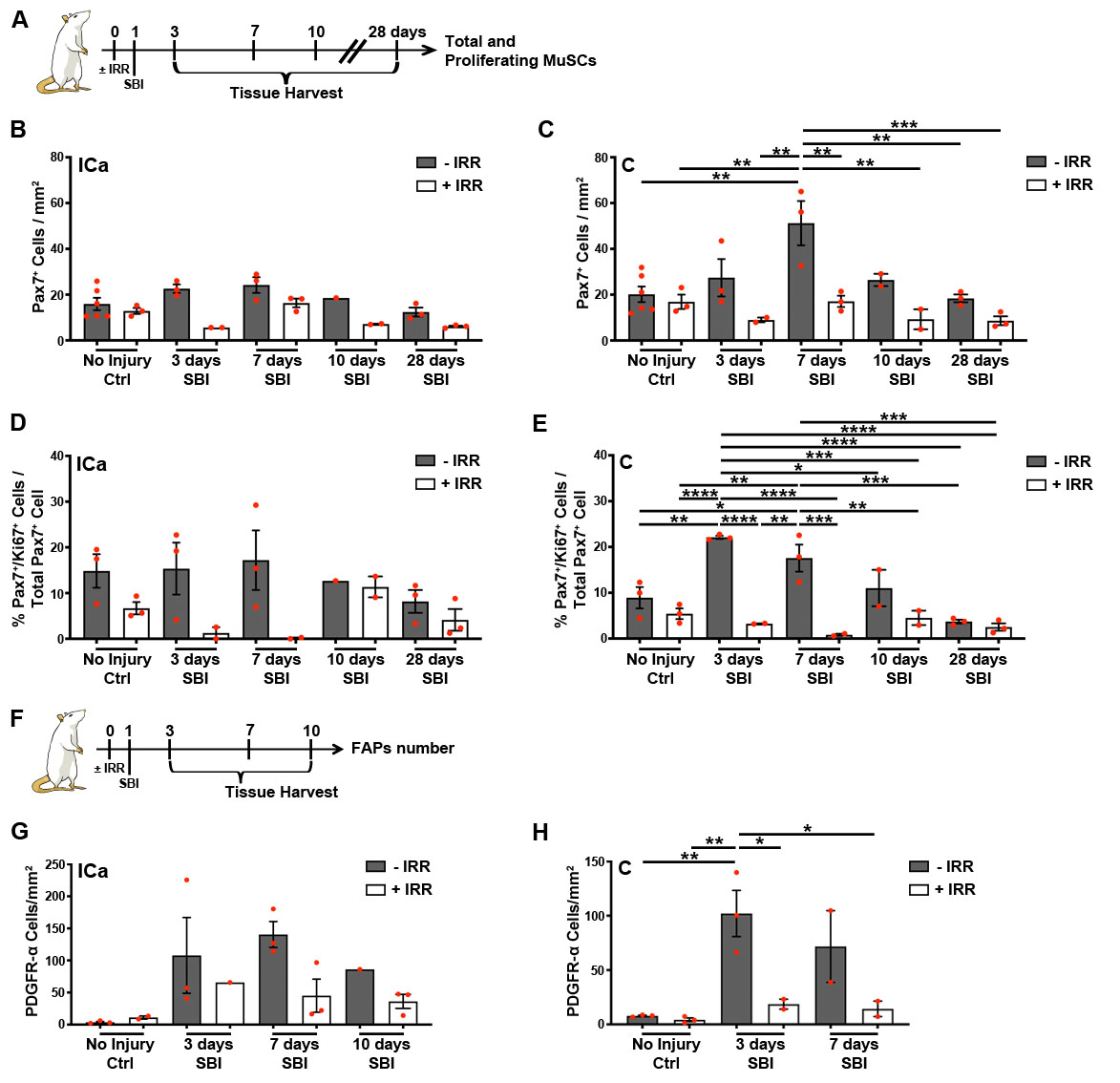

### Supplemental Figure 3

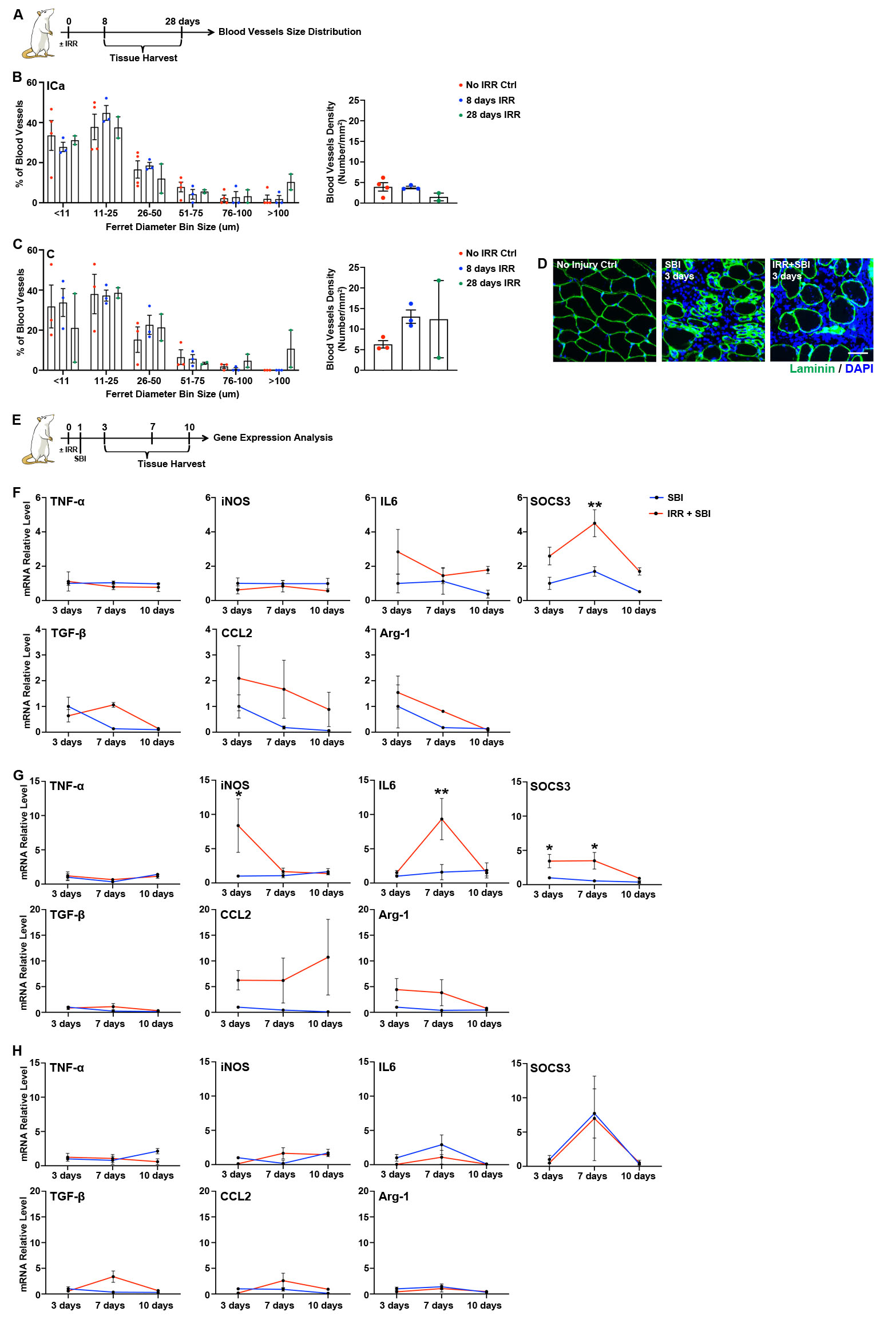

### Supplemental Figure 3

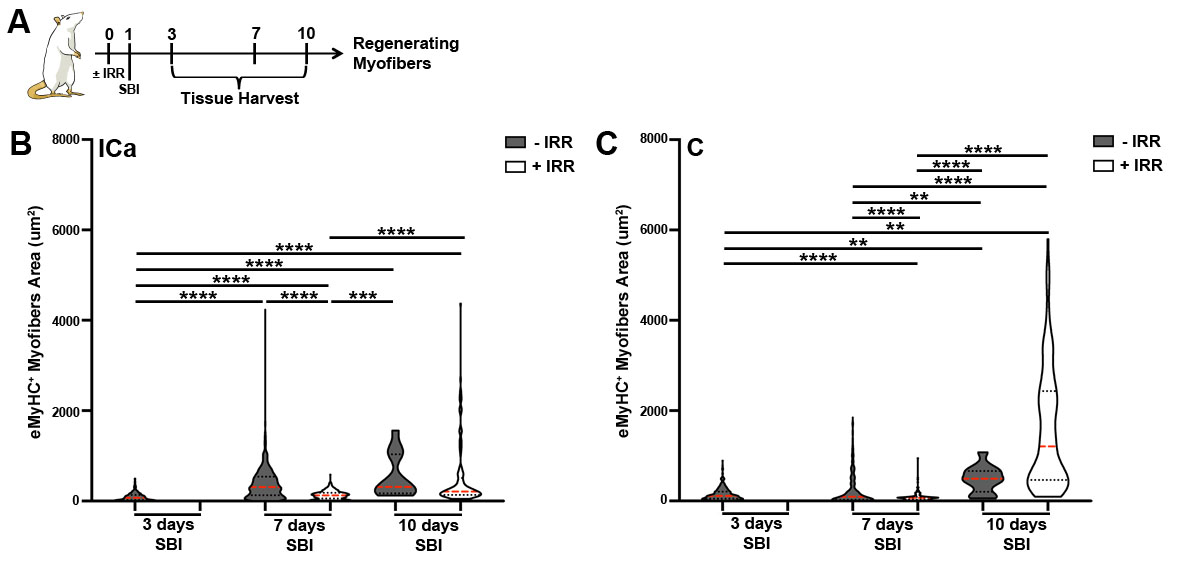
